# A Cell Atlas of Thoracic Aortic Perivascular Adipose Tissue: a focus on mechanotransducers

**DOI:** 10.1101/2023.10.09.561581

**Authors:** Janice M. Thompson, Stephanie W. Watts, Leah Terrian, G. Andres Contreras, Cheryl Rockwell, C. Javier Rendon, Emma Wabel, Lizabeth Lockwood, Sudin Bhattacharya, Rance Nault

## Abstract

Perivascular adipose tissue (PVAT) is increasingly recognized for its function in mechanotransduction. To examine the cell-specificity of recognized mechanotransducers we used single nuclei RNA sequencing (snRNAseq) of the thoracic aorta PVAT (taPVAT) from male Dahl SS rats compared to subscapular brown adipose tissue (BAT). Approximately 30,000 nuclei from taPVAT and BAT each were characterized by snRNAseq, identifying 8 major cell types expected and one unexpected (nuclei with oligodendrocyte marker genes). Cell-specific differential gene expression analysis between taPVAT and BAT identified up to 511 genes (adipocytes) with many (≥20%) being unique to individual cell types. *Piezo1* was the most highly, widely expressed mechanotransducer. Presence of PIEZO1 in the PVAT was confirmed by RNAscope® and IHC; antagonism of PIEZO1 impaired the PVAT’s ability to hold tension. Collectively, the cell compositions of taPVAT and BAT are highly similar, and PIEZO1 is likely a mechanotransducer in taPVAT.

## INTRODUCTION

Perivascular Adipose Tissue (PVAT) is a complex and understudied tissue that surrounds almost all blood vessels in the human body. A seminal finding in 1991 by Soltis and Cassis^1^ changed the view that PVAT served simply as structural support for the vasculature. Specifically, PVAT was recognized to uptake norepinephrine.^2^ Further studies identified that PVAT produces contractile and relaxant substances that affect vascular tone, important findings given that this function changes in cardiovascular disease.^3^ However, this view of PVAT as solely important for secretion of vasoactive factors is limited. For example, PVAT assists in stress relaxation, a form of mechanical support to the artery.^4^ Here, mechanical support of the artery is considered a new function of PVAT. However, the mechanotransducers involved and their expression in the constituent cells of PVAT are not known. As illustrated by this example, there are significant gaps in our understanding of the tissue’s cellular make up and molecular mechanisms. We are committed to the discovery and understanding of new functions of PVAT, such as mechanotransduction. More specifically, here we interrogate PVAT for cellular contributions of mechanotransduction, an action informed by creation of a cell atlas of the Dahl SS rat thoracic aortic PVAT (taPVAT).

This study aims to fill in those gaps by building a comprehensive single-cell atlas of taPVAT using single-nuclei RNA sequencing (snRNAseq) to investigate the cellular distribution of mechanotransducers in PVAT, and contrast it to subscapular brown adipose tissue (BAT), a non-PVAT adipose tissue of the same type.^5^ We focus our study on PVAT from the thoracic aorta because the cellular constituents and type of adipocytes in PVAT are location dependent.^6^ Rat taPVAT most resembles brown adipose tissue, abdominal aorta PVAT resembles both brown and white adipose tissue, and mesenteric PVAT mostly resembles white adipose tissue.^5,7,8^ Bulk microarray studies have found few differences in global gene expression profiles between thoracic PVAT and interscapular BAT in mouse.^5^ However, this difference has not been examined at the single-cell level, which can reveal cell-to-cell heterogeneity commonly masked by bulk gene expression measures.

The thoracic aorta does not share its PVAT with other vessels, as occurs in abdominal vessels, such that conclusions can be made with respect to aortic function confidently. The taPVAT also consist of an anterior strip and two lateral strips (right and left) which are attached to the spine emerging from different developmental origin.^9^ The present snRNAseq experiments allow for construction of a cell atlas, giving insight to the cells that populate this complex tissue. Moreover, we can compare findings from taPVAT to that of BAT from the same rat to determine whether PVAT possesses cell types/genes different from those in BAT. If so, this knowledge could be used to consider potential PVAT specific therapeutic interventions. Our work was carried out in tissues from the Dahl salt sensitive (Dahl SS) rat. The rat and this specific strain of rat are particularly important in the study of hypertensive cardiovascular disease. We identify cell types within taPVAT that express mechanotransducers and test whether one of them, *Piezo1*, participates functionally in mechanotransduction of taPVAT.

## RESULTS

### Characterization of adipose tissue cell populations

To characterize the cell types of the anterior taPVAT and BAT, we used single-nuclei RNA sequencing (snRNAseq) as outlined in Figure 1 (see STAR Methods). A total of 29,703 (taPVAT) and 28,387 (BAT) nuclei passed quality control and ambient RNA removal using SoupX^10^ for taPVAT and BAT, respectively (Figure 2A-B). The median number of unique genes and unique molecular identifier (transcripts), respectively, identified in each nucleus were 1,393 and 2,029 for taPVAT and 1,638 and 2,398 for BAT. Leiden clustering found 20 clusters (Figure S1) which were manually annotated based on marker genes and comparison to previously published adipose tissue datasets^11–14^ identifying 8 major cell types, each represented in similar proportions except for mesothelial cells primarily being present in taPVAT (0.4%, *119* nuclei) compared to BAT (0.004%; *1* nucleus) (Figure 2C).

**Figure 1.**
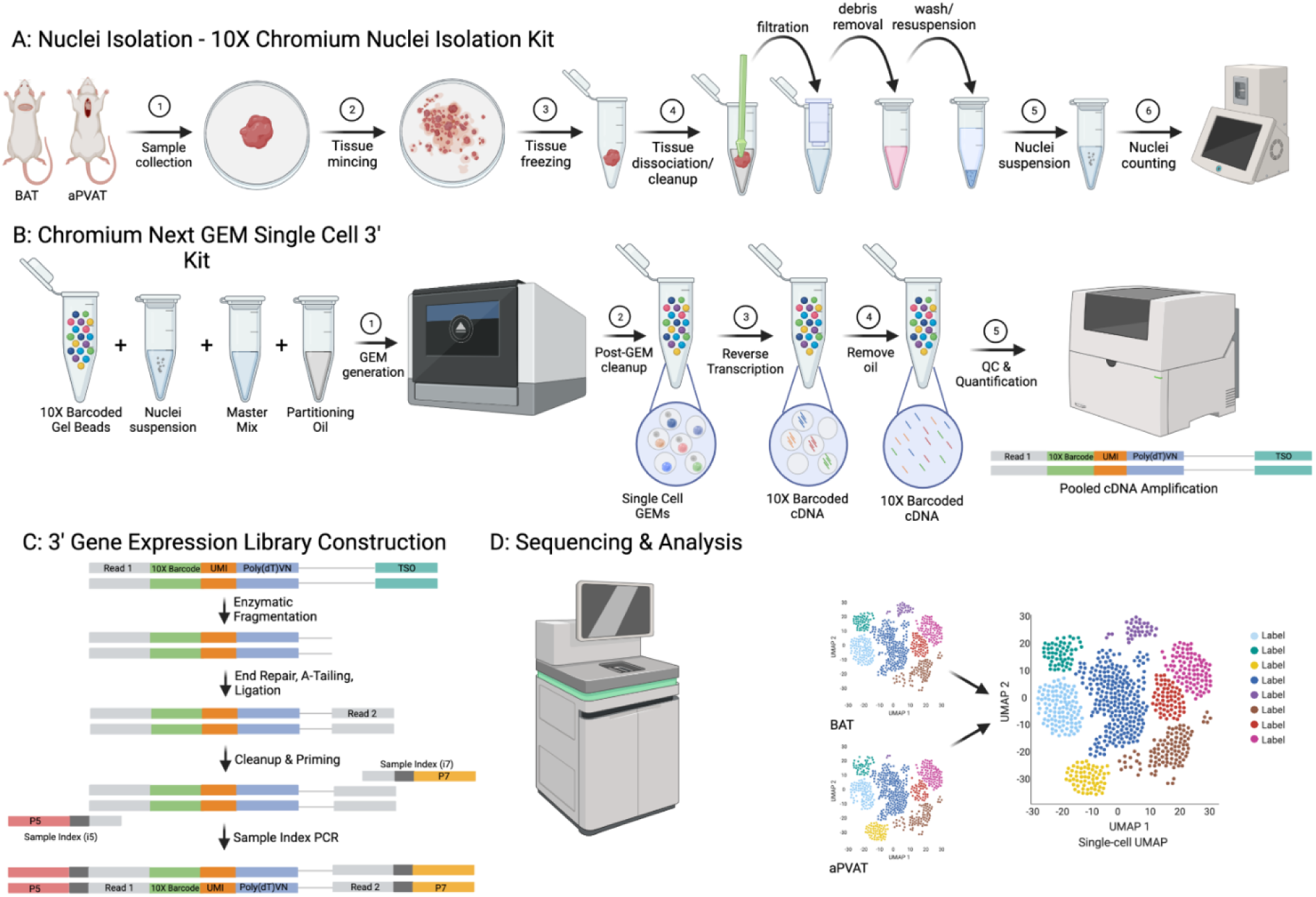
Procedure overview used to derive transcript data to inform cell-based clusters. (A) Nuclear isolation from taPVAT and BAT from male Dahl SS rat. (B) Use of Chromium Next GEM Single Cell 3’ kit to produce amplified cDNA. (C) Construction of a 3’ gene expression library. (D) Sequencing and analyses of cell-based clusters (see STAR Methods for more details). All panels of the workflow overview figure were created using BioRender and some parts of the diagram were modified from the 10X Chromium Nuclei Isolation Kit and Chromium Next GEM Single Cell 3’ Kits user guide.

**Figure 2.**
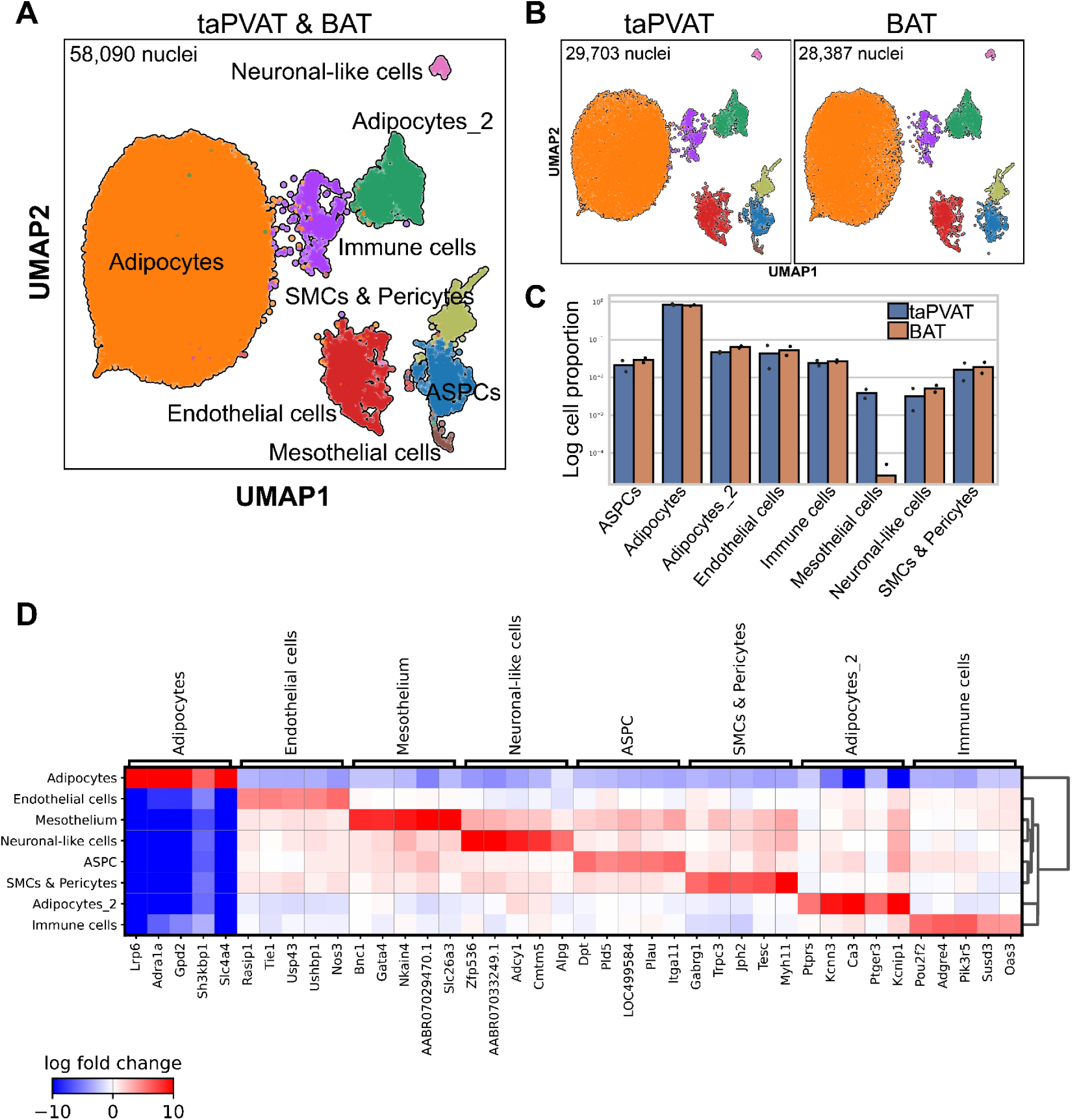
Characterization of the male Dahl SS rat taPVAT and BAT adipose tissue depots. (A) UMAP visualization of nuclei from the taPVAT and BAT following integration using scVI and manual annotation of cell types. (B) UMAP visualization for nuclei from each individual adipose tissue depot. (C) Relative proportion of each cell type for each adipose tissue depot. Proportions are shown on a log scale for comparison of high abundance and low abundance cell types. Bars represent the mean values while individual points represent the proportions in individual samples. (D) Top 5 marker genes for each cell type (listed at top of diagram) identified common to both taPVAT and BAT.

The largest clusters (Leiden clusters 0 – 5, 8 – 13, and 17) were collectively annotated as adipocytes for broad comparison, though clear subtypes are present (*e.g.*, *Acaca* and *Acly* enriched cluster; Figure S2). As expected, adipocytes represent ≥ 80% of captured nuclei in both taPVAT and BAT (Figure 2C). The remaining clusters were annotated according to the highest gene module score for previously identified marker genes in mouse and human white adipose tissue,^13^ or human BAT,^14^ as well as examination of the top marker genes in each Leiden cluster (Figure 2D, S1). Endothelial cells (EC) clearly resembled those of other published datasets and were marked by *Rasip1* which is involved in the formation of EC junctions.^15^ The immune cell cluster is represented by markers of several immune cell types including monocytes, natural killer (NK) cells, T cells, and B cells (Figures S1, S3). Smooth muscle cells (SMCs) and pericytes clustered together, marked by *Myh11* which was previously used to trace the differentiation of SMCs and SMC-like cells into beige adipocytes.^16^ In agreement, SMCs and pericytes clustered closely to adipocyte stem and progenitor cells (ASPCs). One population, mesothelial cells, were only identified in taPVAT. A distinct population of adipocytes was also identified expressing several signaling markers often found in neural cells (e.g., *Ptprs*, *Kcnn3*, and *Ca3*) but clearly resembling adipocytes. Another cluster also sharing several markers with neural cells such as oligodendrocytes and Schwann cells (*Zfp536* and *Sox10*) could not be found in other published adipose tissue datasets.

### BAT and taPVAT cell types are highly similar but exhibit tissue depot-specific gene expression

To examine whether individual cell types of taPVAT closely resemble the cell types in BAT, differential expression (DE) analysis was performed (see STAR Methods). Collectively, the number of DE genes ranged between 552 (Adipocytes) and 16 (neuronal-like cells) across all cell types except for mesothelial cells which could not be compared due to their absence in BAT (Figure 3A). Most DE genes were cell type-specific from 372 in Adipocytes to 39 in SMCs & pericytes while immune cells and neuronal-like cells had ≤ 11 cell type specific DE genes. Consequently, despite the overall similarities between taPVAT and BAT adipocytes, there may be some subtle functional differences associated with their localization in the Dahl SS rat. Closer examination of the 5 DE genes in each cell type with the largest fold-change for taPVAT enriched (induced) and BAT enriched (repressed) individually shows that the biggest differences are represented by only 47 genes (Figure 3B). Among taPVAT enriched genes are lipid metabolism related genes including *Scd* and *Acaca*. Gene set enrichment analysis (GSEA) was used to determine the top 15 taPVAT and BAT cell-specific functions (Figure 3C). While BAT seems to be most enriched in transcriptional regulation as well as cholesterol and carbohydrate metabolism, taPVAT was largely enriched in ion transport, signaling, and chemokine functions. These findings may point to taPVAT playing a greater role in signaling in response to cues from the thoracic aorta.

**Figure 3.**
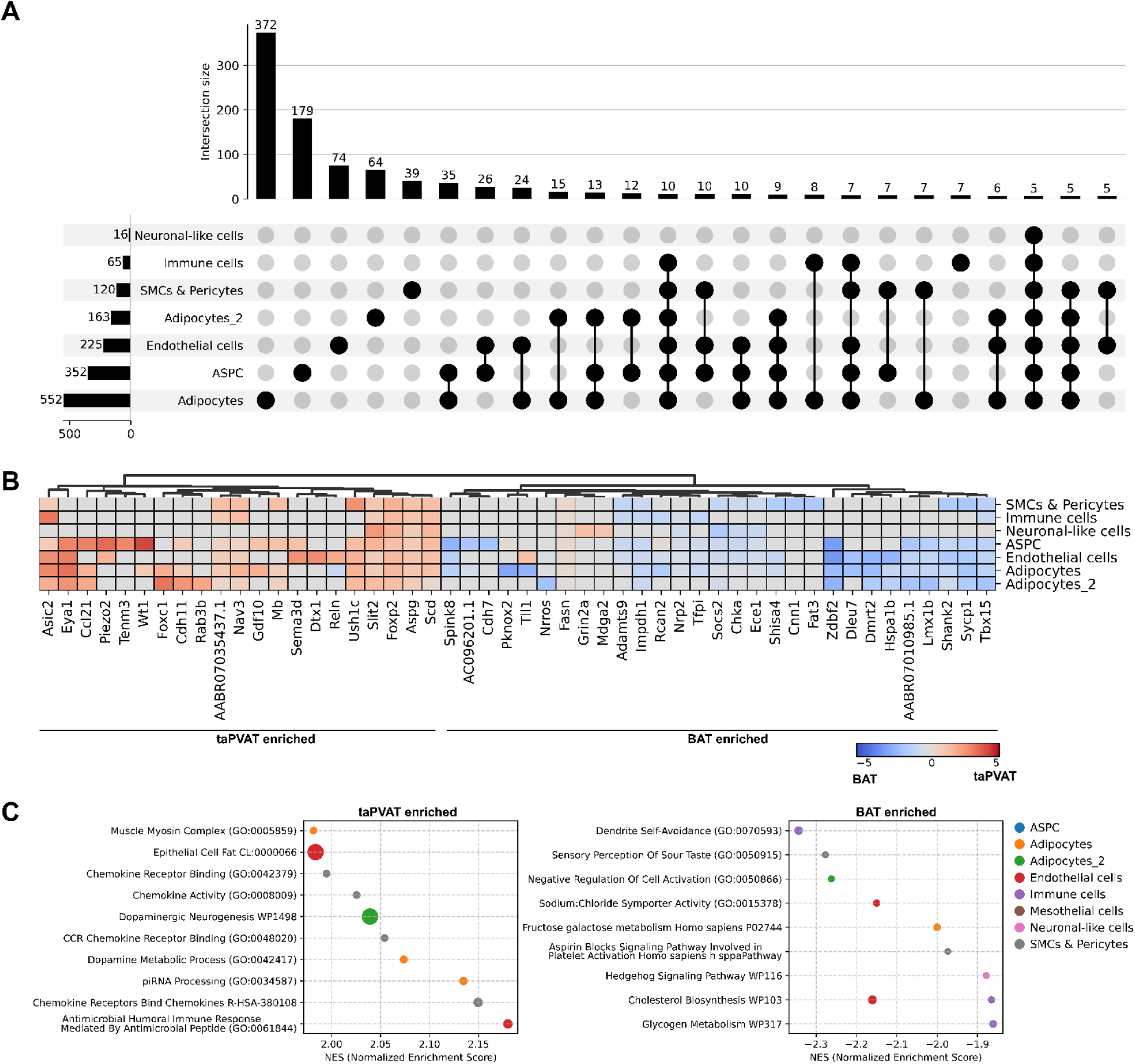
Comparison of taPVAT and BAT adipose tissue depot gene expression. (A) UpSet plot of differentially expressed genes (|fold-change| ≥ 2, adjusted p-value ≤ 0.05) for individual cell types between taPVAT and BAT. The total number of differentially expressed genes for each cell type is shown as horizontal bars on the left and the intersecting list of differentially expressed genes for the cell types markers by a black dot is shown as vertical bar. (B) Heatmap of the top 5 taPVAT and BAT enriched genes. (C) Top 10 enriched functional groups determined using GSEApy (see STAR Methods) where a positive NES (top) represents taPVAT enriched functions and negative NES (bottom) represents BAT enriched functions.

### PIEZO1 is a functional mechanotransducer in taPVAT

We use snRNAseq to characterize cell populations in two adipose tissue depots considered to be similar in phenotype (*i.e.,* brown fat) but which are expected to be at least somewhat functionally distinct due to taPVAT being constantly exposed to mechanical forces from blood flow in the aorta, the largest conduit vessel. Therefore, we examined the expression of mechanotransduction genes in both adipose tissue depots (Figure 4A). Rat genes functionally associated with mechanosensing were identified in Gene Ontology representing 3 broad categories; mechanosensitive ion channels, mechanosensitive cation channels, and genes implicated in sensing mechanical stimuli. Known but currently unassigned genes implicated in mechanosensing such as *Ddr2* and *Itgb1* were also included because of their actions as receptors for collagen. GSEA of each cell type found that mesothelial cells were enriched in genes implicated in the detection of mechanical stimuli while the adipocyte_2 cluster as well as endothelial cells were enriched in mechanosensitive ion/cation channels (Figure 4B). Among the mechanosensing genes showing the largest fold-change between taPVAT and BAT were the acid sensing ion channel *Asic2* and mechanosensitive ion channel *Piezo2* (Figure 4C). However, both *Asic2* and *Piezo2* were lowly expressed across all clusters with *Piezo2* not falling within the top expressed mechanotransduction related genes (Figure 4A). Conversely, some mechanosensing-related genes were expressed in specific cell types and differentially expressed between taPVAT and BAT including *Ddr2* (ASPCs; BAT enriched), *Cxcl12* (endothelial cells; taPVAT enriched), *Ano1* (SMCs and pericytes; BAT enriched), *Fyn* (ASPCs; BAT enriched) (Figures 4A, D). Two mechanosensing genes were highly expressed homogenously. *Piezo1* was found in all cell types and *Slc12a2* was highly expressed in all cell types except the adipocyte_2 population.

**Figure 4.**
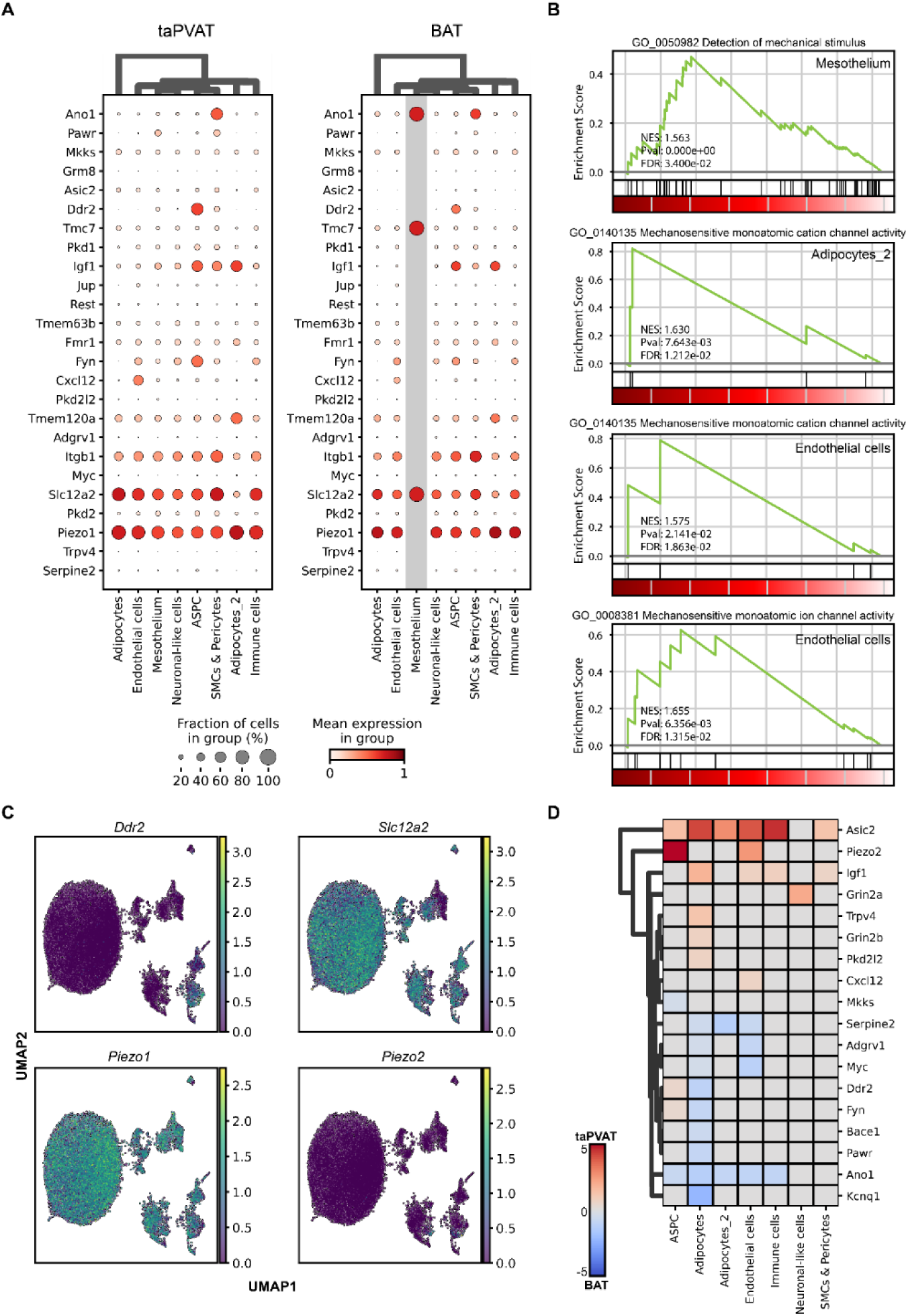
Analysis of mechanotransduction related gene expression. (A) Dot plot of the most highly expressed mechanotransduction genes in taPVAT and BAT. Dot size represents the percent of nuclei expressing the gene and the color intensity represents the expression level. The mesothelium of BAT is shaded in grey to indicate that values may appear higher than expected as only 1 nucleus was present for this tissue. (B) Cell-specific enrichment of mechanotransduction related GO terms (see STAR Methods). (C) UMAP visualization of expression for *Ddr2*, *Slc12a2*, *Piezo1*, and *Piezo2*. (D) Heatmap of fold-change for differentially expressed mechanotransduction related genes (|fold-change| ≥ 2, adjusted p-value ≤ 0.05).

To test the hypothesis that PIEZO1 is functionally important in taPVAT, we first used RNAScope® to validate the presence of *Piezo1* mRNA in the PVAT (anterior and lateral) and minimal expression in the media of the isolated thoracic aorta from the male Dahl SS rat (Figure 5A). The same pattern of expression can be seen in female Dahl SS rats (Figure S4). Similarly, PIEZO1 protein, identified immunohistochemically, was present in the taPVAT (anterior and lateral) but not media of the thoracic aorta; these sections were taken from the same aorta as was used for RNAscope® experiments (Figure 5B). The functional importance of PIEZO1 was studied in the isolated PVAT ring and aortic ring from the Dahl SS male rat, measuring isometric tension as the primary outcome. The PIEZO1 antagonist GsMTx4 caused a profound loss of ability of the tissue to hold tension during stress relaxation in the isolated PVAT but not in the aorta (Figure 5C). Collectively, these findings are consistent with the snRNAseq data in locating *Piezo1* mRNA to taPVAT, with PIEZO1 playing a functional role mediating the response to the mechanical intervention of stretch.

**Figure 5.**
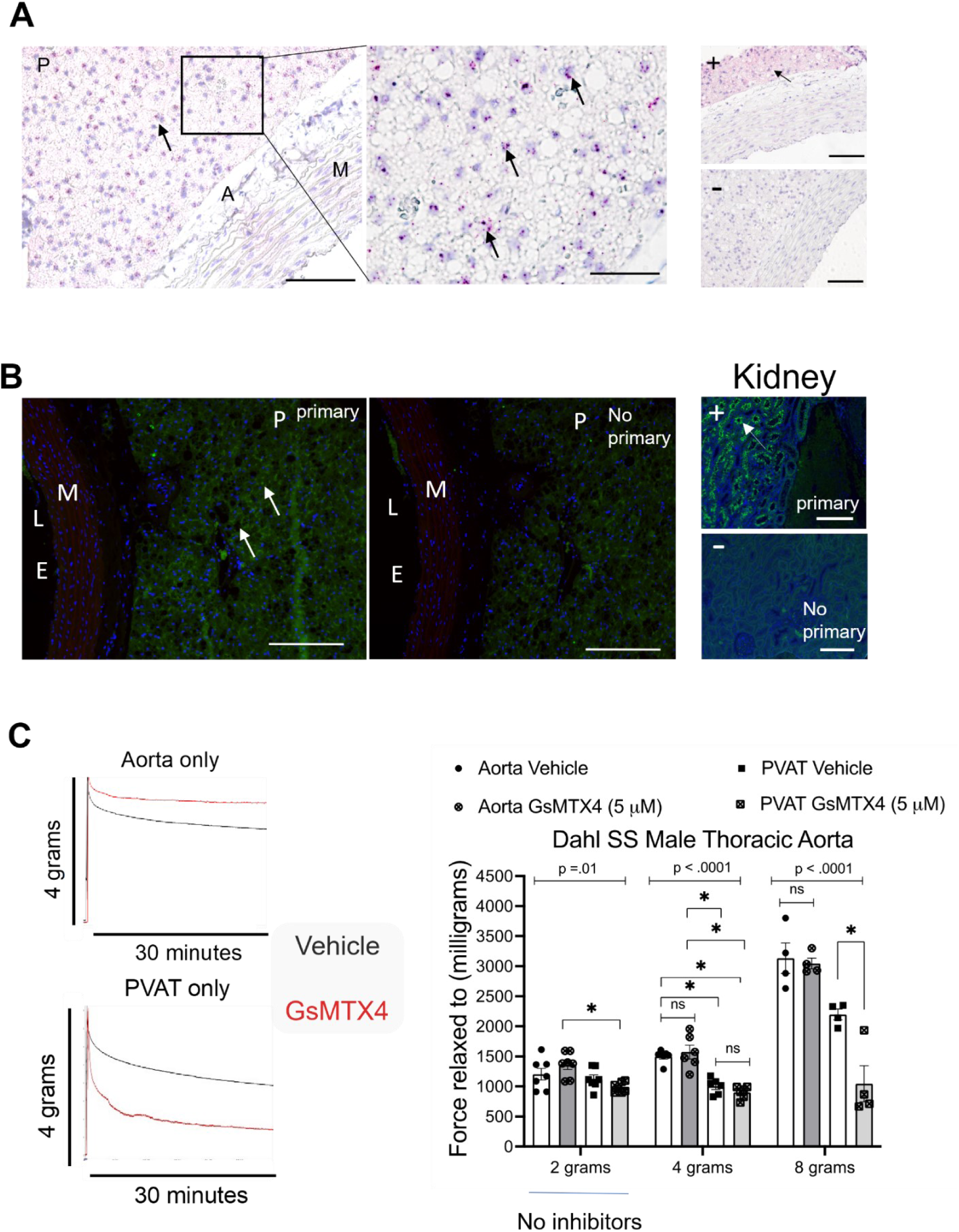
The mechanotransducer Piezo1 is expressed in and functionally serves PVAT of the thoracic aorta. (A) Brightfield of *Piezo1* mRNA ZZ-probe detection (RNAScope) in the thoracic aorta of the Dahl SS male rat. Arrows indicate positive dot detection. P = PVAT (anterior), A = adventitia, M = Media. Horizontal bar indicates 50 micrometers. To the right are tissue-specific positive (+) controls for housekeeping gene Peptidylprolyl isomerase B (PPIB) and negative (-) controls for bacterial gene Diaminopimelate (DapB). Images are representative of three male rats. (B) Fluorescent immunohistochemical detection of PVAT. Images of sections incubated with (1^st^ full image) and without primary (2^nd^ full image) PIEZO1 antibody. P = PVAT (anterior), M = Media, L = Lumen, E = Endothelium. Far right images are of the kidney nephron tubule epithelium staining for Piezo1 in the presence (top) and absence (bottom) of PIEZO1 primary antibody. Representative of three male rats. (C) Left: Representative tracing of the response of the aorta (top) or its surrounding ring of PVAT (bottom) to a 4 gram passive tension addition in the absence (black) or presence of PIEZO1 inhibitor GsMTx4 (5 μM). Right: Quantifies tension relaxed to after application of a baseline tension of 2 grams (no inhibitors present); after addition of 2 grams tension to achieve 4 grams total (after 1 hour incubation with vehicle/inhibitor); and after addition of 4 grams tension to achieve 8 grams total (after reincubation with vehicle/inhibitor). Bars are means ± SEM with individual scattered values. Asterisk (*) marks statistically significant (p<0.05) differences within group members as marked by lines.

### ASPC subpopulations express different levels of *Piezo1*

The ASPCs are reported to consist of multiple subpopulations with distinct functional roles.^17–19^ As expected, these cells expressed elevated levels of common markers including *Dcn*, *Fbn1*, *Cd34*, and *Pdgfra* (Figure 6A). We re-integrated and clustered the ASPCs which were represented by 744 nuclei in taPVAT and 770 nuclei in BAT, resulting in 3 Leiden clusters (Figure 6B). Among the top markers was *Pi16* (cluster 1) previously found to represent a non-proliferating population,^18^ while *Bmper*, elevated in cells which differentiate into brown adipocytes,^18^ was high in cluster 2. The proportion of cells did not widely differ between taPVAT and BAT (Figure S5). More importantly, when evaluating the expression of *Piezo1* in these subpopulations, it was found to be primarily expressed in *Pi16*^high^ ASPCs and much lower in *Bmper*^high^ nuclei (Figure 6D).

**Figure 6.**
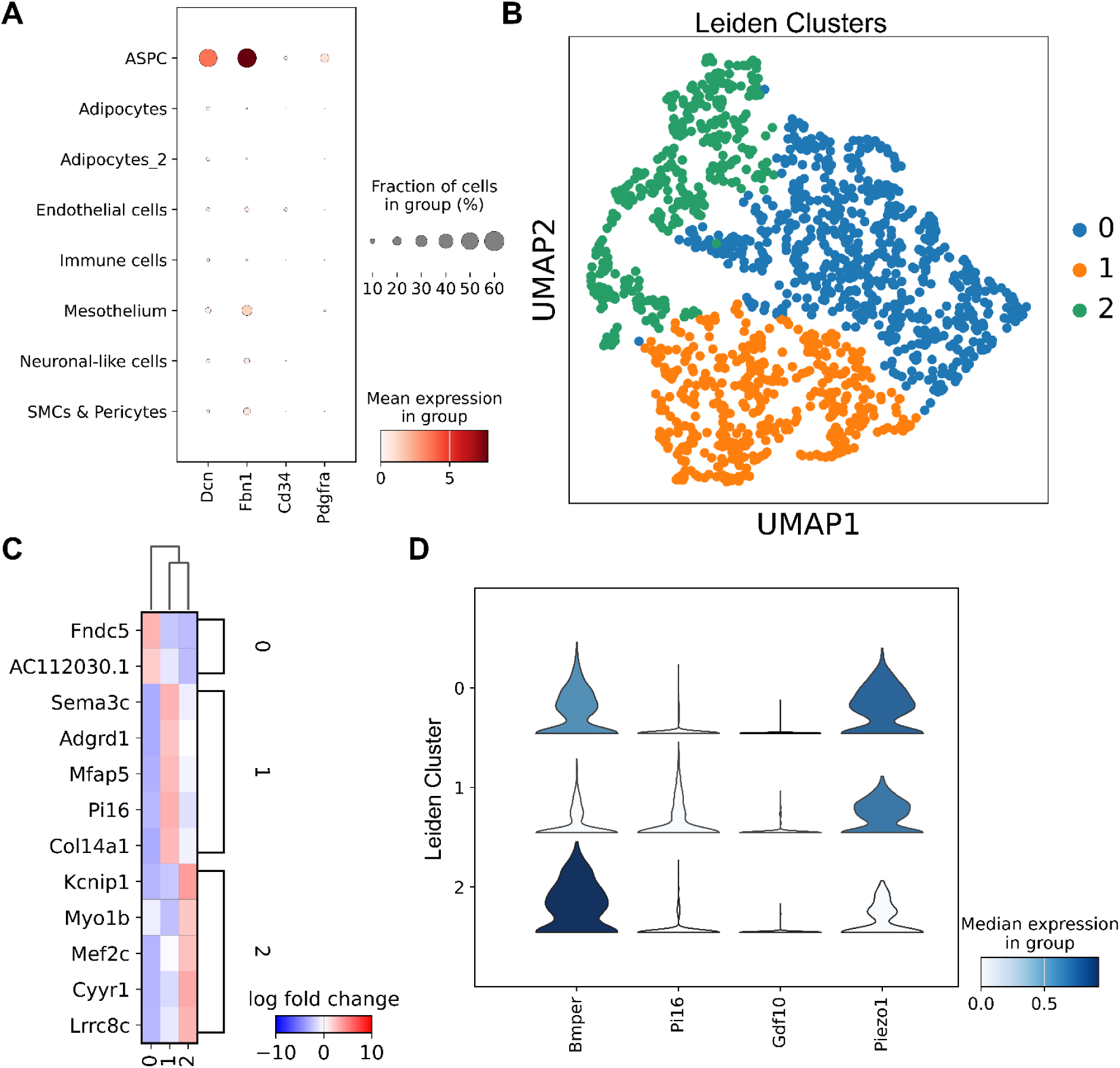
Adipocyte stem and progenitor cells subtypes in PVAT from Dahl SS. (A) Expression dot plot of ASPC marker genes *Dcn*, *Fbn1*, *Cd34*, and *Pdgfra* in individual cell types for taPVAT and BAT combined. The dot size represents the percent of genes expressing the gene while the color represents the mean expression. (B) UMAP visualization of re-integrated ASPCs as described in STAR Method. Nuclei were re-clustered using Leiden clustering at a resolution of 0.1 identifying a total of 3 subpopulations. (C) Top 5 markers genes for each ASPC Leiden cluster with a minimum |fold-change| of 2. Only 2 genes met the threshold criteria for cluster 0. (D) Median expression and distribution of expression for previously identified markers *Bmper*, *Pi16*, and *Gdf10* are shown as violin plots for each Leiden cluster identified in panel B. *Piezo1* expression is also shown for each Leiden cluster.

## DISCUSSION

### Cell Atlas of the rat taPVAT and BAT

To our knowledge, this work is the first cell atlas of a PVAT in a rat, one specifically important to the field of hypertension. Eight major cell types were represented in taPVAT: two adipocyte subtypes, endothelial cells, immune cells, neuronal-like cells; SMCs and pericytes, mesothelial cells, and ASPCs. The presence of most cells types was expected, and that of immune cells is consistent with previous work that identified T cells, B cells, macrophages, neutrophils and mast cells in taPVAT of the Dahl SS male and female rat.^20^ While this study did not break down the immune cell cluster for more comprehensive analysis, further sub-clustering (Figure S3) shows that monocytes, T cells, and B cells were all present in both taPVAT and BAT. Smooth muscle and endothelial cells are likely from the microvasculature of the taPVAT as we ensured that experimental samples contained no *tunica media*. Two cell type clusters could not be clearly matched to previously identified mouse and human datasets; adipocytes_2 and neuronal-like cells. The presence of neuronal cells in taPVAT is controversial; no study has demonstrated definitive innervation of adipocytes in PVAT. Analysis of single-cell thoracic aorta between E18 and P3 identified a population of Schwann cells marked by the expression of *Mpz* which were not present in adult samples.^17^ Our data showed low expression of *Mpz* but significant expression of several other markers found in Schwann cells and oligodendrocytes.^17,21^ Adipocytes_2 were notable in their expression of the ion channel *Kcnn3* which may be reflective of stem-cell like characteristics.^22^

### Similarities of taPVAT and BAT

Previous bulk gene expression analysis of taPVAT and BAT in mice has shown that both brown adipose tissue depots are highly similar.^5^ Consistent with these mice, our data identified genes enriched in PVAT, *Cfh* and *C7*, both immunomodulatory genes. Similarly, the BAT enriched gene *Tbx15*, a putative master regulator of adipose tissue genes,^23^ was identified. However, the total number of DE genes was comparatively greater in our study. This might be attributed to the difference in species, the use of next generation sequencing technology, and/or the ability to examine cell-specific differences. Nevertheless, snRNAseq of the two brown adipose tissue depots in Dahl SS rats show strong similarities in cellular composition and overall gene expression. Interestingly, both taPVAT and BAT expressed many of the same mechanotransducers at comparable levels including *Piezo1* and *Slc12a2* (*aka* NKCC1). It was not the goal of this work to understand differences in mechanotransducer expression between BAT and taPVAT. Nonetheless, it is notable that BAT expressed many of the same mechanotransducer transcripts found in taPVAT. It remains unclear why BAT would express mechanotransducers, though evidence points to a link between stiffness of brown adipocytes and their thermogenic function.^24^

### Dahl SS rat is an important model for CVD

The Dahl SS rat has been a mainstay model in hypertension research since its original discovery by Lewis Dahl.^25^ This rat strain, fed a high (4% or above) salt diet, develops elevated blood pressure. The Medical College of Wisconsin (MCW) has led important efforts to understand this rat genetically in the hopes of identifying a gene/genes responsible for hypertension.^26^ Importantly, both researchers at MCW and MSU find that a high fat (HF, 60%kCal, lard) diet, independent of salt, creates a hypertension in the Dahl SS when fed the diet from weaning.^27,28^ The HF-diet fed hypertension occurs in both males and females, making this a valuable model. It is for these reasons that the Dahl SS rat was our choice of model in the present study. Our work is also important because a vast majority of cell work in adipose tissues (of all kinds) have been done in the mouse and human and not the rat. This work thus lays a foundational study for an adipose tissue cell atlas in a rat.

### Piezo1 as a functional mechanotransducer in PVAT

SnRNA seq identified multiple recognized mechanotransducers in taPVAT of the male Dahl SS rat. While not exhaustive, major classes of mechanotransducers were interrogated. We focused on membrane limited mechanotransducers (ion, cation channels). Of these gene ontologies, *Piezo1*^29^ was the most highly and homogenously expressed in cells populating taPVAT. Consistent with this finding, mRNA for *Piezo1* was observed in taPVAT, as was protein. Importantly, inhibition of PIEZO1 function by GsMTx4^30^ caused a loss of the ability of taPVAT to maintain tone after a stretch challenge. GsMTx4 differs from Dooku1, a more widely used antagonist, in that GsMTx4 can inhibit the endogenous function of PIEZO1. By contrast, Dooku1 inhibits those actions stimulated by the agonist Yoda1.^31^ Notably, neither RNAscope® nor immunohistochemistry detected *Piezo1* mRNA or protein, respectively, in the media of the same vessel. This lack of measurable *Piezo1* explains the overall lack of effect of GsMTx4 on tone of the isolated aorta. PVAT thus becomes important as a site for PIEZO1’s role in aortic function. This includes the clinical measure of aortic stiffness as measured by pulse wave velocity, with elevated aortic stiffness serving as an independent risk factor for cardiovascular disease.^32^ The independent roles, however, of this mechanosensor within each subtype of ASPC in which *Piezo1* was identified remains unknown. We recognize that we have not delved into the many other mechanotransducers that could function in PVAT, including the focal adhesion kinases, cytoskeletal proteins, and other adhesion receptors.^33^ Specifically, the present data support future investigation of *Slc12a2* (Na K Cl transporter, NKCC1) in PVAT. Nonetheless, using *Piezo1*, the present work exemplifies a way in which snRNAseq and physiological/pharmacological studies can be complementary.

Consistent with previous studies in other adipose tissue depots identifying ASPC subpopulations,^17–19^ this study also found subpopulations expressing similar markers identified in BAT^18^ including *Bmper*^high^ and *Pi16*^high^ populations while *Gdf10* was lowly expressed in all clusters. The presence of the cells in taPVAT demonstrates the similarities between ASPC from taPVAT and BAT. Previous evidence has shown that mechano-activation of PIEZO1 reduces the potential of ASPCs to become adipocytes.^34^ Higher expression of Piezo1 in *Pi16*^high^ ASPCs compared to *Bmper*^high^ ASPCs reflects the differentiation and proliferation potential of each population.^18,34^ The relationship between PIEZO1 and adipogenesis of ASPCs warrants further investigation.

PVAT, especially that around the thoracic aorta, is being recognized as a tissue that works in concert with the rest of the vessel to determine aortic stiffness.^35,36^ Thoracic aortic stiffness, measured mechanically, was lower when PVAT was included.^37^ Advanced glycation end-products^38^ and interleukin-6^39^ both contributed to arterial stiffness in the mouse at least in part from actions within PVAT. Finally, the peroxisome proliferator activated receptor gamma (PPARγ) activator pioglitazone improved PVAT microenvironment of the *ob/ob* mouse to potentially reduce aortic stiffness.^40^ These findings are consistent with the serendipitous discovery in the PPARγ smooth muscle cell specific knockout mouse. In this KO mouse, PVAT did not develop around the thoracic aorta and the KO mouse had a higher pulse wave velocity, a surrogate of aortic stiffness, than the WT.^41^ All together, these findings support that PVAT – now known to possess mechanotransducers – should be considered mechanistically in the determination of arterial stiffness.

### Limitations of the study

We recognize experimental limitations of the present work. The anterior strip of PVAT was used for snRNAseq. The lateral strips, those which hold the aorta to the spine, were not included in the snRNA seq analyses but were present in all remaining experiments. Lineage tracing in the mouse found the anterior PVAT adipocytes were derived from smooth muscle 22α^+^ (SM22α^+^) progenitors, while the lateral PVAT possessed both SM22α^+^ and Myf5^+^ cells.^9^ It is unknown whether this different developmental program occurs in the rat. Along these lines, reference datasets and functional annotations were derived primarily from mouse/human resources given that these resources are limited in the rat. Tissues from males were the focus given the intention to understand whether this study was feasible. Similarly, the present work studied brown/brown-like adipose tissue, as opposed to white, given that the PVAT around the thoracic aorta is brown/brown-like. Future experiments in females and white PVAT fat (mesenteric) will be important to carry out. We also recognize the limitations of using single nuclei vs single cell bias.^42^ Nuclei vs cell isolation was a deliberate choice given the fragility of isolated adipocytes and difficulty of working in a high lipid environment, as well as with buoyant, large cells. SnRNAseq provided an effective means to determine the cell types that populate the PVAT of the thoracic aorta.

### Conclusions

The taPVAT and BAT of the male Dahl SS rat are comprised of eight dominant cell types. The chief cell type difference between these two brown fats was a greater representation of mesothelial cells in taPVAT vs BAT. In taPVAT, most cell types possessed mechanotransducer genes, with *Piezo1* and *Slc12a2* being the most highly represented transcripts. Using isolated taPVAT, PIEZO1 was shown to be functionally important for maintaining tone in the stretch-challenged tissue. Collectively, this work defines the cells that reside in PVAT; that these are more similar than different to those in BAT; and that cells of taPVAT possess (functional) mechanotransducers that contribute to the discovery of new functions of this tissue. Two goals were fulfilled with this study. First, we now have a good sense of the major types of cells in taPVAT. Such knowledge helps us better understand the potential physiology of this tissue. Second, we now know that taPVAT is more similar to BAT in its cellular and gene composition than it is different. This provides a challenge in developing PVAT specific therapies based on the cell or gene.

## ACKNOWLEDGEMENTS

This work was supported by the National Heart, Lung, and Blood Institute [NHLBI P01HL152951] to JT, SWW, LT, GAC, CR, CJR, LL, SB, RN. EW was supported by T32 [T32GM142521] to the Department of Pharmacology and Toxicology at Michigan State University.

## AUTHOR CONTRIBUTIONS

**CRediT: JT** - Formal analysis, Investigation, Data Curation, Writing - Original Draft; Visualization**; SWW** - Conceptualization, Formal Analysis, Investigation, Resources, Data curation, Writing - Review & Editing, Supervision, Project administration, Funding acquisition; **LT** - Software, Formal analysis, Investigation, Data Curation, Writing - Review & Editing; Visualization; **GAC**-Validation, Supervision, Writing - Review & Editing; **CR** - Validation, Supervision, Writing - Review & Editing; **CJR** - Formal analyis, investigation, Writing-Review & Editing, Visualization; **EW** - Formal analysis, Investigation, Data Curation, Writing - Review & Editing, Visualization; **LL** - Formal analysis, Investigation, Data Curation, Writing - Review & Editing, Visualization; **SB** - Writing - Review & Editing, Supervision, Funding acquisition; **RN** - Methodology, Resources, Writing - Review & Editing, Supervision, Funding acquisition.

## STAR★METHODS (Extended)

### KEY RESOURCES TABLE

| REAGENT or RESOURCE | SOURCE | IDENTIFIER |
| --- | --- | --- |
| **Antibodies** |  |  |
| Piezo1 | Alomone Laboratories | APC-087 |
| Alexafluor 488 goat anti – rabbit IgG | Invitrogen | A11008 |
| **Chemicals, peptides, and recombinant protein** |  |  |
| Fatal Plus | Vortech Pharmaceuticals | V.P.L. 9373 |
| 10% Bovine Serum Albumin in DPBS | Millipore-Sigma | A1595-50mL |
| Sodium chloride | VWR | BDH9286 |
| Potassium chloride | Avantor | 3040-05 |
| Potassium phosphate, monobasic | Sigma-Aldrich | 795488 |
| Magnesium sulfate heptahydrate | VWR | 0662-500G |
| Sodium bicarbonate | Sigma-Aldrich | S6014 |
| dextrose | Sigma-Aldrich | D9434 |
| EDTA, disodium salt, dihydrate | J.T. Baker | 8993-01 |
| Calcium chloride | Sigma-Aldrich | C7902 |
| phenylephrine | Sigma-Aldrich | P6126 |
| Phosphate-Buffered Saline without Calcium and Magnesium | Millipore-Sigma | D8537-500ML |
| Molecular Grade Nuclease-Free Water | Invitrogen | AM9930 |
| ViaStain AOPI Staining Solution | Nexcelom | CS2-0106-5ml |
| Low TE Buffer | Thermo Fisher Scientific | 2502246 |
| Buffer EB | Qiagen | 1014609 |
| 10% Tween 20 | Bio-Rad | 1662404 |
| SPRIselect | Beckman Coulter | 19686100 |
| Trizma hydrochloride | Sigma-Aldrich | T5941 |
| Tween-20 | Sigma-Aldrich | P1379 |
| GsMTx4 | MedChemExpress | HY-P1410 |
| Histochoice Clearing Agent | VWR | H103-4L |
| Isopropanol | VWR | 9084-03 |
| Antigen Unmasking Solution | Vector Laboratories | H3301 |
| Normal Goat Serum | Vector Laboratories | S-1000 |
| Dulbecco's Phosphate Buffered Saline | Sigma- Aldrich | D8537 |
| Antifade Mounting Medium with DAPI | Vector Laboratories | H-1500 |
| 10% formalin | Fisher | SF100-4 |
| Xylenes | Fisher Chemical | X3P-1GAL |
| 100% Ethanol | VWR | 89125-170 |
| Hydrogen peroxide | ACDBio | 322330 |
| Target Retrieval Reagent | ACDBio | 322000 |
| Protease Plus | ACDBio | 322330 |
| hematoxylin | American MasterTech Scientific | HXGHE1LT |
| ammonia | Honeywell Fluka | 221228 |
| Ecomount | BioCare Medical | EM897L |
| Commercial assays and kits |  |  |
| 10X Genomics Nuclei Isolation Kit | 10X Genomics | PN-1000494 |
| Chromium Single Cell 3' Reagent Kit v3.1 | 10X Genomics | PN-1000269 |
| D5000 ScreenTape | Agilent |  |
| D1000 ScreenTape | Agilent |  |
| KAPA Library Quantification Kit | Roche | 07960255001 |
| RNAScope RED | ACDBio | 322350 |
| Rat Piezo1 ZZ probes | ACDBio | 587591 |
| Peptidylpropyl isomerase B (Ppid) probe | ACDBio | 313921 |
| Diaminopimelate (DapB) probe | ACDBio | 310043 |
| Deposited data |  |  |
| Next Generation Sequencing Data | This study | GSE244451, SCP2384 |
| Software and algorithms |  |  |
| CellRanger 7.1.0 | 43 | support.10xgenomics.com/single-cell-gene-expression/software/downloads/latest |
| scvi-tools_0.20.0 | 44 | github.com/scverse/scvi-tools |
| scanpy 1.9.1 | 45 | github.com/scverse/scanpy |
| scDblFinder 1.14.0 | 46 | github.com/plger/scDblFinder |
| SoupX 1.6.2 | 10 | github.com/constantAmateur/SoupX |
| Seurat 4.3.0.1 | 47 | github.com/satijalab/seurat |
| Python 3.9.12 | https://www.python.org/ | www.python.org/ |
| R 4.3.1 | R Foundation | www.r-project.org/ |
| GSEAPy 1.0.5 | 48 | github.com/zqfang/GSEAPy |
| PyDESeq2 0.4.0 | 49 | github.com/owkin/PyDESeq2 |
| Ensembl Rat Reference Genome (Rnor_6.0) | 50 | Assembly: GCA_000001895.4 |
| TapeStation Analysis Software 4.1 | Agilent | https://www.agilent.com |
| Prism 10.0.01 | GraphPad | https://www.graphpad.com |
| LabChart 8.1.25 | ADInstrument | https://www.adinstruments.com |
| NIS Elements | Nikon | BR.`4.60 64 bit |
| KAPA Library Quantification Data Analysis Template | Roche |  |
| BioRender | BioRender | www.biorender.com |
| Other |  |  |
| Agilent TapeStation 4200 | Agilent | G2991BA |
| NovaSeq 6000 | Illumina |  |
| Cellometer Vision | Nexcelom |  |
| Chromium controller | 10X Genomics | 1000204 |
| SimpliAmp ThermalCycler | Applied Biosystems | A24812 |
| QuantStudio 6 Flex Real-Time PCR System | Applied Biosystems |  |
| Nikon Microscope | Nikon | Nikon Eclipse Ti-S |
| Nikon Camera | Nikon | DS-Qi1MC |
| Isometric transducers | Grass Instruments | FT03 |
| PowerLabC with Octet Bridge | ADInstruments |  |

### RESOURCE AVAILABILITY

#### Lead contact

Further information and requests for resources should be directed to and will be fulfilled by the lead contact, Dr. Rance Nault.

#### Materials availability

This study did not generate new unique reagents.

#### Data and code availability

Raw and processed snRNAseq data are publicly available on the Gene Expression Omnibus (GEO; https://www.ncbi.nlm.nih.gov/geo/) at accession ID GSE244451 and for visualization and querying on the Broad Single-Cell Portal (SCP; https://portals.broadinstitute.org/single_cell) with the accession id SCP2384. Analysis code is available on Github (https://github.com/naultran/BAT_taPVAT_snRNAseq).

## METHOD DETAILS

### EXPERIMENTAL MODEL AND STUDY PARTICIPANT DETAILS

#### Animal models and tissue collection

Dahl SS male rats were obtained from a colony at MSU bred from parents purchased from Charles River Laboratory (provided samples for snRNA seq experiments) or purchased from Charles River (RNAscope®, immunohistochemistry, and isometric contraction analyses). Animals were on a normal diet [Teklad 22/5 Rodent diet (Madison WI, USA)] and housed in a 12:12 light:dark light cycle. Food and drinking water were available *ad libitum*. Procedures using animals complied with National Institutes of Health Guide for the Care and Use of Laboratory Animals (2011) and approved by the MSU Institutional Animal Care and Use Committee (PROTO202000009). This study was conducted with Animal Research: Reporting of *In Vivo* Experiments (ARRIVE) guidelines (essential 10 and recommended) in mind.^51^

Before tissue removal, rats were given pentobarbital as a deep anesthetic (80 mg kg^-1^, ip). A bilateral pneumothorax was created prior to vessel dissection. Tissues for snRNAseq experiments [thoracic aorta PVAT (taPVAT) and subscapular brown adipose tissue (BAT)] were dissected as whole pieces from each animal. For taPVAT, the anterior strip of taPVAT (not attached to spine) was used for snRNA seq experiments. BAT was removed between the scapula. The taPVAT was dissected from the vessel while BAT was cleaned of adherent white fat (subcutaneous) under a stereomicroscope and in a Silastic®-coated dish filled with physiological salt solution [PSS in mM: NaCl 130; KCl 4.7; KH_2_PO_4_ 1.18; MgSO_4_ • 7H_2_O 1.17; NaHCO_3_ 14.8; dextrose 5.5; CaNa_2_EDTA 0.03, CaCl_2_ 1.6 (pH 7.2)]. A 50 mg piece of each taPVAT and BAT was minced prior to snap freezing in liquid nitrogen and storage at –80 °C. For RNAScope® experiments, aorta with complete PVAT and liver were cleaned of blood and formalin (10%)-fixed for 16-32 hours, then embedded in paraffin blocks. These same tissues were used for fluorescent microscopy. For isometric contractility experiments, the thoracic aorta was dissected from the aortic arch to the diaphragm and placed in PSS.

#### Nuclei isolation and single-nuclei RNA sequencing

Nuclei from frozen minced aortic perivascular adipose tissue or subscapular brown adipose tissue (∼ 50 mg) were isolated with the Chromium Nuclei Isolation Kit (10X Genomics) according to manufacturer’s recommended protocol (CG000505, RevA), with a final resuspension in 50 µL Wash and Resuspension Buffer. Following isolation, nuclei concentration was determined with a Nexcelom Cellometer Vision and Viastain AOPI staining solution. The 10X Chromium Next GEM Single Cell 3’ Reagent Kit v 3.1 (Dual Index) was used with a targeted cell recovery of 10,000 according to manufacturer’s recommended protocol (CG000315, Rev D). Following GEM generation, barcoding, cleanup and amplification, generated cDNA was quantified, and purity determined with an Agilent 4200 TapeStation using a D5000 ScreenTape assay. To generate the single nuclei 3’ gene expression library, 25% of the total cDNA obtained underwent fragmentation, end repair, A-tailing, and adaptor ligation, followed by cleanup with SPRIselect reagent. Individual 10X Sample Index names from the Dual Index Plate TT Set A were recorded and added to the appropriate sample, then amplified according to manufacturer’s protocol. A final cleanup with SPRIselect reagent was done, with samples eluted into 35.5µL Buffer EB. The average fragment size following library construction was determined with an Agilent 4200 TapeStation using a D1000 ScreenTape assay.

#### Library Quantification

Quantification of library samples was done with the KAPA Library Quantification Kit. Library samples were diluted (1:100, 1:1000, 1:5000, 1:10,000) in DNA Dilution Buffer (10 mM Tris-HCl, pH 8.0 −8.5 + 0.05% Tween 20). In triplicate, 4 µL of each diluted sample and kit supplied DNA Standards were pipetted into a standard 96-well PCR plate containing 6µL KAPA SYBR FAST qPCR Master Mix. No Template Controls containing 4 µL water were run in duplicate. An adhesive film to prevent sample evaporation was placed over the top of the plate and the plate centrifuged at 1000 rpm for 1 minute. The following conditions were run in a QuantStudio 6 Flex Real-Time PCR System: 1 cycle of 95°C 5 minutes; 35 cycles of 95°C 30 seconds, 60°C 45 seconds. A standard melt curve was run to ensure adapter dimers were not present. Calculations were performed in the KAPA Library Quantification Data Analysis Template to determine the undiluted library concentration. Libraries were submitted to Novogene for 150bp paired-end sequencing on a NovaSeq6000 at a target depth of 50,000 reads/nuclei.

#### Quality control and preprocessing

Initial quality control was performed using FastQC. Reads were aligned to the rat reference genome Rnor_6.0 (assembly GCA_000001895.4) using the 10x CellRanger v7.1.0 pipeline.^43^ Scanpy^45^ and other commonly used python packages (*i.e.*, numpy, pandas, matplotlib) were then used to do additional quality control, preprocessing, analyzing, and visualization of the data. Any nuclei that had less than 200 detected genes or genes that were detected in less than 3 cells were discarded. Cells that were found to be expressing greater than 10% mitochondrial genes were also discarded. Ambient RNA and doublet removal was done using SoupX^10^ and scDblFinder^46^ respectively. To reduce technical variation between samples, batch correction with the python package scVI was performed.^44^ Principal component analysis (PCA) and uniform manifold approximation and projection (UMAP) were used to project the high-dimensional data into a 2D space. After dimensionality reduction, Leiden unsupervised clustering method was used to group the cells into distinct subpopulations based on their gene expression patterns.

#### Marker identification, cell annotation, and differential expression

Leiden clustering was performed at resolutions of 0.1 to 1.5 to identify clearly distinct clusters resulting in a total of 20 clusters at a resolution of 1.0. To support cluster annotation, cell type marker lists were obtained from previously published human and mouse white adipose tissue snRNAseq^13^ and human brown adipose tissue scRNAseq.^14^ The marker lists were sorted by fold-change and the top 50 (or all markers when fewer than 50) were used to calculate gene module scores with a control size matching the marker length and a total of 500 bins. Additionally, Wilcoxon Rank Sum test was used to identify the top markers sorted by fold-change relative to all other clusters combined. Collectively, these results were used to manually annotate the clusters based on internal expertise as well as cross-referencing to literature. To simplify cell annotation and differential expression, the largest cluster likely representing subtypes or different cell states of adipocytes were collapsed into an overarching adipocyte annotation. Re-clustering of only ASPC (*Dcn, Fbn1, Pdgfra*) was performed to highlight the differences within the ASCP group.

Recent studies suggest that use of pseudobulk and pseudoreplicates for differential expression analysis of single-cell transcriptomic data can reduce false discoveries.^52,53^ A total of 3 pseudoreplicates for each adipose tissue were randomly sampled from the integrated data and aggregated into individual pseudobulk samples. PyDESeq2^49^ was used to perform differential expression analysis using tissue as experimental design factor for each cell type independently. Genes were considered differentially expressed when the adjusted p-value ≤ 0.05.

#### Gene set enrichment analyses

Rat specific genes implicated in mechanotransduction were identified in the Gene Ontology web tool AmiGO (https://amigo.geneontology.org/amigo) by looking up known mechanotransducers and related terms. A total of three GO terms were identified: GO:0008381 Mechanosensitive monoatomic ion channel activity, GO:0140135 Mechanosensitive monoatomic cation channel activity, GO:0050982 Detection of mechanical stimulus. When gene sets were analyzed collectively, known mechanotransduction genes *Ddr2* and *Itgb1* were also included as they were not present in these lists. Scores for each of these ontologies were calculated as described in the cluster annotation method and shown as dot plots. Gene set enrichment analysis was performed on pre-ranked genes for each cell types, ranked according to expression level, and then analyzed using GSEAPy.^48^

#### RNAScope®

Sections (5 microns thick) from formalin-fixed, paraffin embedded thoracic aorta + PVAT were cut by the MSU Investigative Histopathology laboratory, mounted on Superfrost® Plus microscope slides (cat. # 12-550-15, Thermo Scientific, Waltham, MA, USA), and stored at 4°C after air-drying. To de-paraffinize, slides were baked one hour in a 60°C oven and then treated by a series of washes in xylene and 100% ethanol. Slides were incubated for 10 minutes in hydrogen peroxide (proprietary grade) to permeabilize. RNA retrieval was performed first by boiling slides in the Target Retrieval reagent (cat. #322000; Advanced Cell Diagnostics, Hayward, CA, USA) for 8 minutes and then by incubating slides with Protease Plus solution (cat. #322330, Advanced Cell Diagnostics) for 15 minutes. For *In situ* detection of Piezo1 mRNA, the RNAscope® 2.5 HD Assay-RED kit (cat. #322350, Advanced Cell Diagnostics) according was used according to the manufacturer’s protocol. Slides were incubated with a probe for rat *Piezo1* (cat. #587591, Advanced Cell Diagnostics), the housekeeping gene *Peptidylprolyl isomerase B* (*Ppib*) for a positive control for mRNA detection, or the bacterial gene *Diaminopimelate* (DapB) for an mRNA species that should not be present as a negative control (cat. #313921, #310043, Advanced Cell Diagnostics) for two hours at 40°C. To amplify the probe signal, a series of amplification reagents was run [Amp 1: 30 minutes at 40°C, Amp 2: 15 minutes at 40°C, Amp 3: 30 minutes at 40°C, Amp 4: 15 minutes at 40°C, Amp 5: 30 minutes at RT, Amp 6: 15 minutes at RT, “Red” A+B reagent: 10 minutes at RT (cat. #322360, Advanced Cell Diagnostics)]. Slides were counterstained with hematoxylin for 2 minutes and 0.02% ammonia water for 10 seconds at room temperature then mounted with EcoMount (cat. #EM897L, BioCare Medical, Pacheco, CA) and a glass coverslip. Images were taken with a Nikon Digital Sight DS-Qil camera and Nikon NIS Elements BR 4.6 software on a Nikon TE2000 inverted microscope. All background correction was applied to the full image and was consistent across positive and negative controls and images were brightened or contrasted as a whole, never in part. Positive staining was determined by red punctate dots in the cell.

#### Immunofluorescence

Sections (5 microns thick) from formalin-fixed, paraffin embedded Dahl SS rat thoracic aorta + PVAT and kidney (positive control for PIEZO1) were cut by the MSU Investigative Histopathology laboratory, mounted on Superfrost® Plus microscope slides (Thermo Scientific,12-550-15) and were air dried and stored at room temperature (RT). To de-paraffinize, slides were washed 2 times with Histochoice Clearing Agent (VWR, H103) and 4 times with isopropanol (VWR,9084-03) and 2 times with distilled water for 3 minutes (min) each wash. Antigen retrieval was performed by boiling slides for 30 seconds in Antigen Unmasking Solution (Vector Laboratories, H3301). Slides were rinsed in distilled water and air dried. To contain the blocking serum, primary and secondary antibody solutions, circles were drawn around the sections with an ImmEdge Hydrophobic pen (Vector Laboratories, H-4000). Slides were incubated at RT with 1.5% normal goat serum (Vector Laboratories, S-1000) in Dulbecco’s phosphate buffered saline (PBS) (Sigma-Aldrich, D8537) blocking solution (BS) for 1 hour (hr). Positive control sections were incubated with 1:600 primary antibody Anti-Piezo1 (Alomone Labs, APC-087) in 1.5% normal goat serum BS and negative control sections were incubated with BS overnight at 4°C. The primary antibody and BS were removed from the sections and the slides were rinsed in Dulbecco’s PBS 3 times for 5 min each rinse. The slides were incubated in 1:1000 secondary antibody AlexaFluor 488 (Invitrogen, ThermoFisher, A11008) for 1 hr at RT. The secondary antibody was removed, and slides were rinsed 3 times with Dulbecco’s PBS for 5 min and allowed to dry thoroughly at RT. Vectashield with DAPI (Vector Laboratories, H-1500) was applied to the sections and cover slips mounted to the slides. The slides were allowed to dry and harden at RT. Images were acquired at 360, 488 and 544nm on a Nikon Eclipse T*i*-S microscope, using a 10x objective, Nikon DS-Qi1MC camera, and NIS elements BR 4.6 software.

#### Isometric Contractility

For creating separated rings of the thoracic aorta and its surrounding PVAT, a section of the thoracic aorta + PVAT was stood on its end in the silastic dish filled with PSS. Two insect pins were inserted into the lumen of the aorta to hold the vessel open. While gently holding the PVAT layer away from the adventitia with fine forceps, small vannas scissors were used to sever connections around the circumference of the vessel between the PVAT and vessel/adventitia. The ring was flipped vertically, and this process repeated. The separated ring of PVAT was then lifted off the vessel. All tissues [PVAT alone or aorta alone] were placed onto two L-shaped stainless-steel rings. Rings were mounted in warmed (37°C) and aerated (95% O_2_, 5% CO_2_) tissue baths (10 ml volume) on Grass isometric transducers (FT03; Grass instruments, Quincy, MA, USA) connected to an 8 channel PowerLab C through an Octet Bridge (ADInstruments, Colorado Springs, CO, USA). Sample type (aortic or PVAT ring) and exposure to vehicle or inhibitor were randomized daily in one of eight different tissue baths. Care was taken to apply no tension on the tissue prior to initiation of the experiment.

All rings started at a tension of 0 grams, equilibrating in the warm buffer for one hour prior to experimentation. During this hour, buffer was exchanged every 15 minutes. Aortic rings were first challenged with a passive tension application of 2 grams applied over ∼15 seconds, applied through clockwise turning of a rack and pinion. Tissues were allowed to relax to this stretch for 30 minutes; the tension achieved at the end of this period recorded. Tissues were then challenged with a maximum concentration of phenylephrine (PE, 10^-5^ M). This response plateaued and was recorded. Tissues were washed identically and repeatedly through 30 minutes to achieve a stable baseline. This first 2 gram challenge was considered a viability test. Next, tissues were incubated for one hour with either vehicle (water) or the PIEZO1 antagonist GsMTx4 (5 μM). Another 2 grams of passive tension (for a total of 4 grams) were applied at the end of this hour, tissues relaxed over 30 minutes and the tension relaxed to recorded. Tissues were challenged with PE (10^-5^ M) again and washed to baseline. A subset of tissues was then reincubated with vehicle/GsMTx4 for 45 minutes without washing. A 4-gram passive tension was applied (total to 8 grams). Thirty minutes, with no washing, were allotted for tissues to relax and tension relaxed to recorded. Tissues were again challenged with PE. Tissues were weighed once the experiment was completed.Lab Chart 8.1.25 (ADInstruments, Colorodo Springs, CO, USA) was used to capture tissue tension/responses through a Mac Mini computer connected to a monitor. Lab Chart allows for capture of actual tracings shown as well as quantitative responses. The magnitude of the response relaxed to at the end of the 30-minute period was recorded for the responses at 2 grams, 4 grams total applied and 8 grams total applied. Data are reported as mean ± SEM for the number of tissues reported. Data are graphed in GraphPad Prism 9. A one-way ANOVA was used at each magnitude of total stretch applied (2, 4, 8) to determine statistical differences between groups. This test, using Bartletts, also verified statistically equivalent variances in groups. * indicates a difference of < 0.05 and brackets point out the specific differences identified by a Tukey’s *post-hoc* test.

## SUPPLEMENTAL INFORMATION

**Figure S1.**
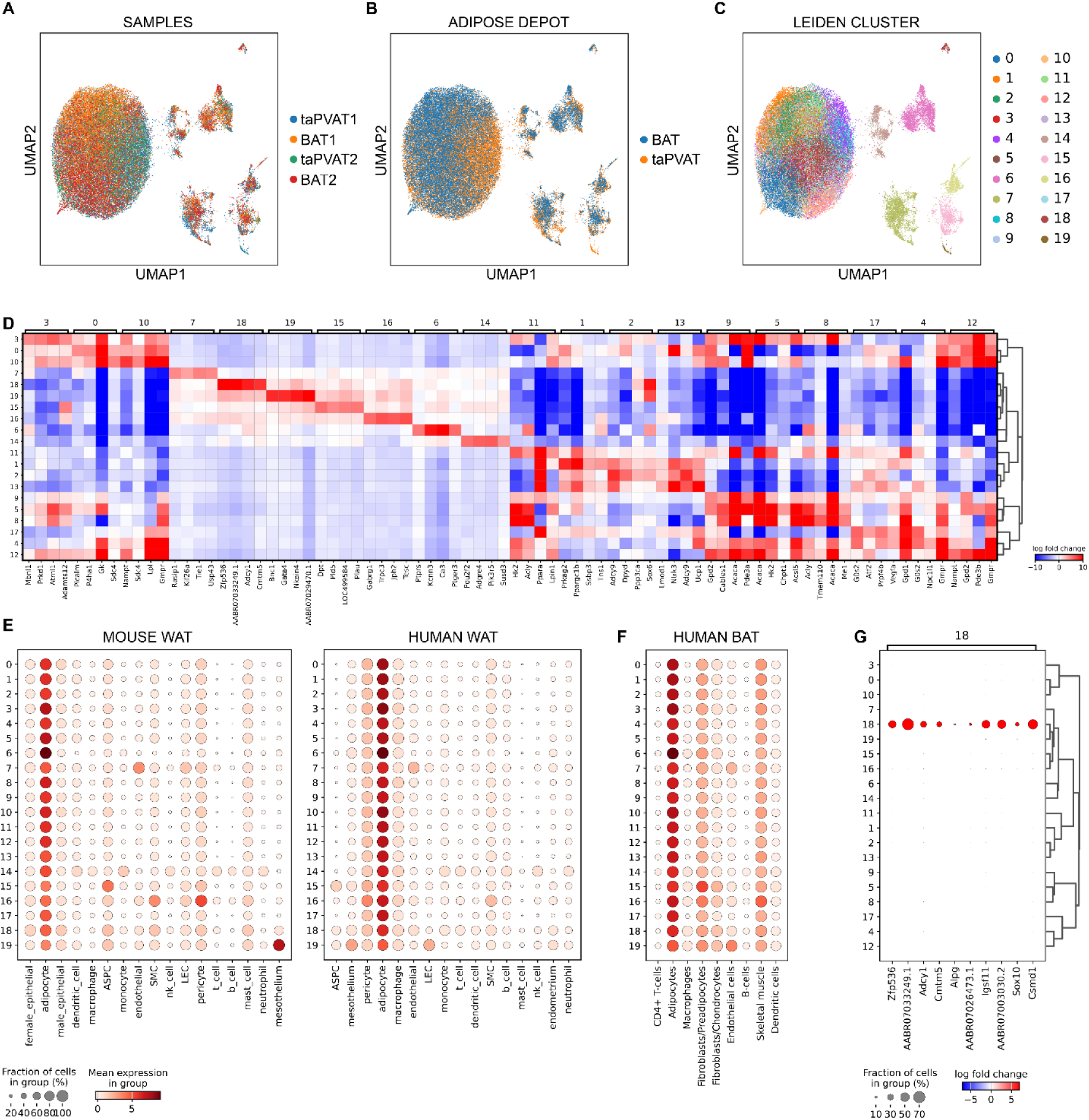
UMAP clustering and cell annotation of Leiden clusters. UMAP visualization of integrated taPVAT and BAT snRNAseq data grouped by (A) sample, (B) adipose tissue source, and (C) Leiden cluster at a resolution of 1.0 chosen to best capture distinct clusters. (D) Heatmap of the top 4 differentially expressed genes (markers) between individual Leiden clusters and all other clusters. (E) Gene module scores for each Leiden cluster using previously identified top 50 mouse and human white adipose tissue (WAT) marker genes (Emont *et al.*, 2022) as gene module. (F) Gene module scores for each cluster using top 50 previously identified BAT cell type markers (Sun *et al.*, 2020). (G) Marker genes for Leiden cluster 18 which was not clearly identified using examined previously published markers.

**Figure S2.**
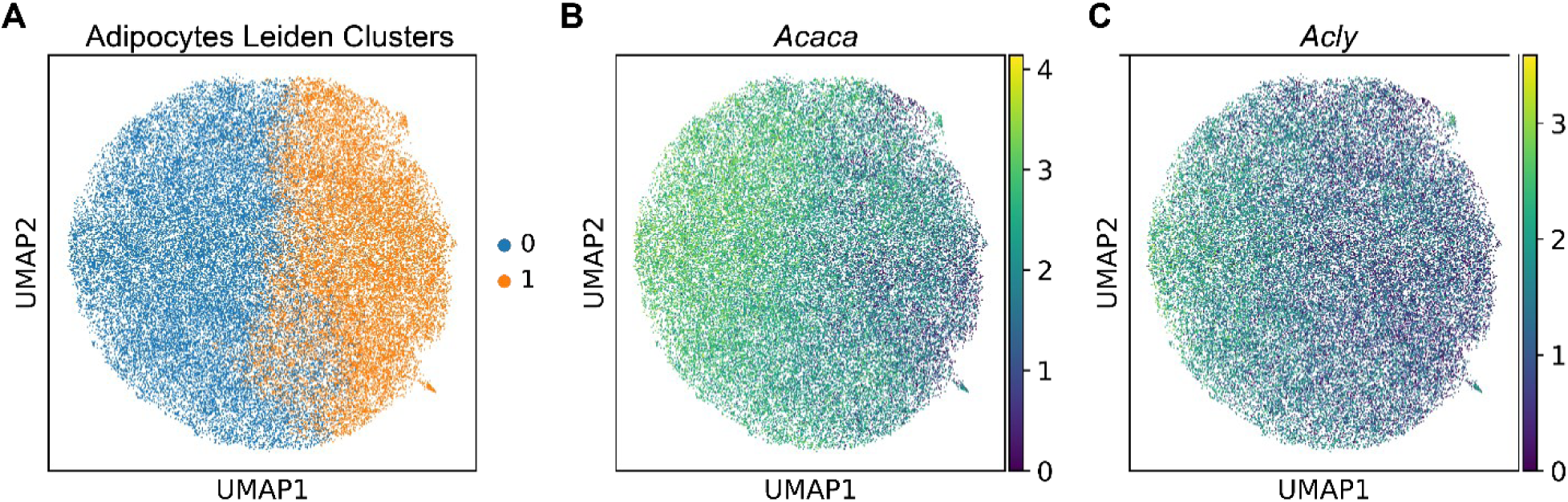
Re-integration and clustering of adipocytes. The adipocyte cluster was extracted from the full snRNAseq dataset, reintegrated, and reclustered to identify putative sub-populations. Two potential sub-populations were identified.

**Figure S3.**
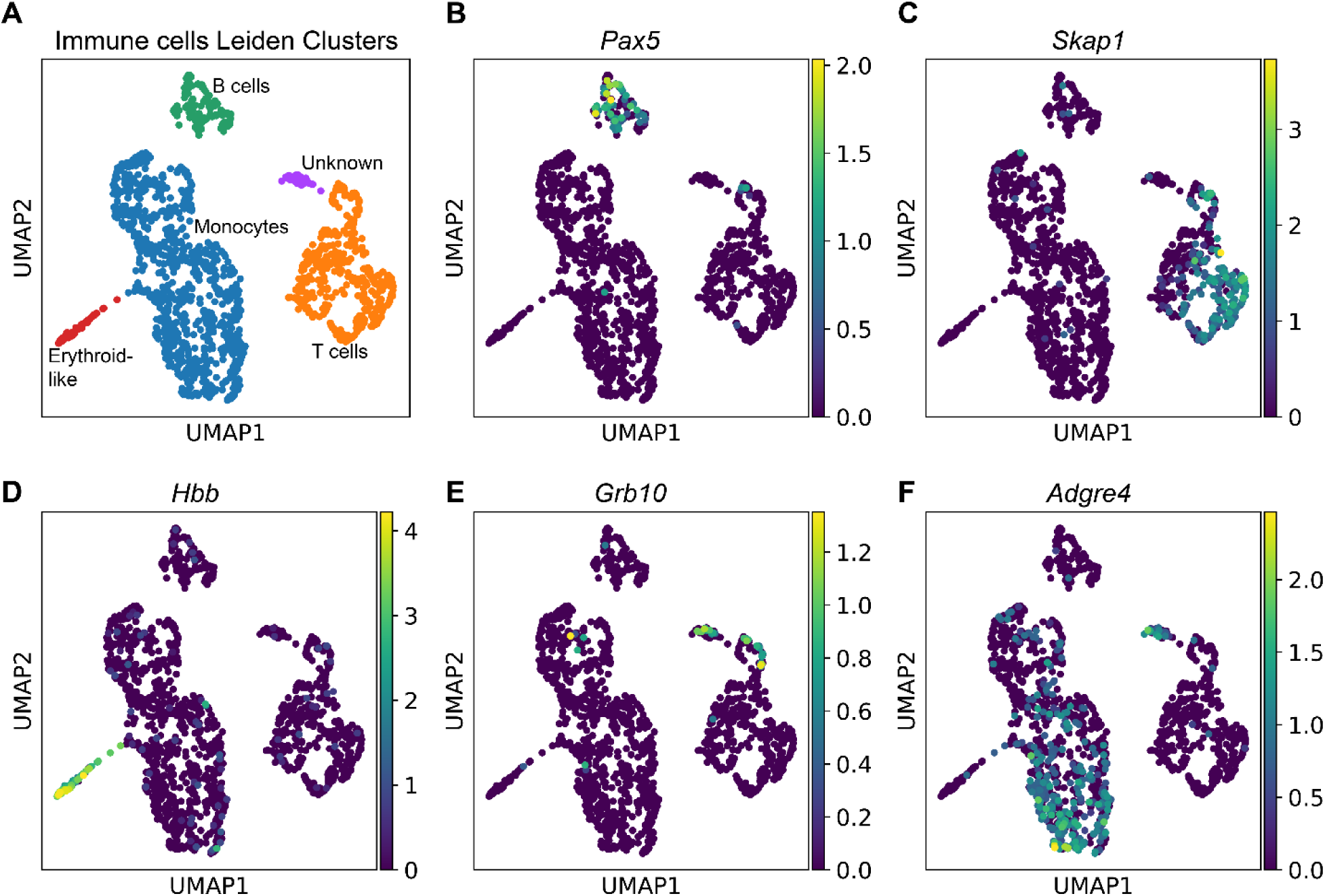
Re-integration and clustering of immune cells. The immune cells cluster was extracted from the full snRNAseq dataset, reintegrated, and reclustered to identify putative sub-populations. Five sub-populations were identified representing erythroid-like; monocytes, B cells, T cells, and another yet to be identified population.

**Figure S4.**
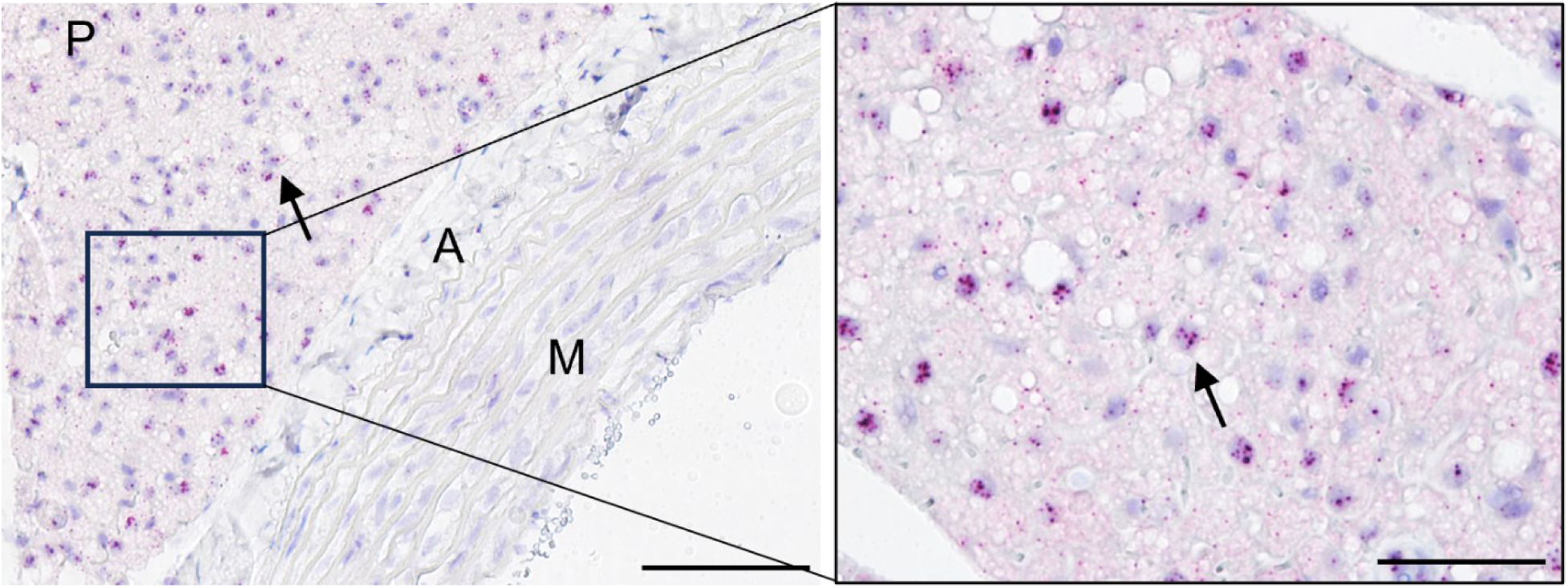
Piezo1 RNAScope analysis of taPVAT from female Dahl SS rats. RNAScope® single plex chromogenic representation of *Piezo1* in Dahl SS female thoracic aorta perivascular adipose tissue (P) [minimal signal detected in adventitia (A), and media/intima (M)]. Each red punctate dot denotes a *Piezo1* transcript. Images are magnified 20x and representative of at least 5 different Dahl SS female rats. Scale bar = 100μm.

**Figure S5.**
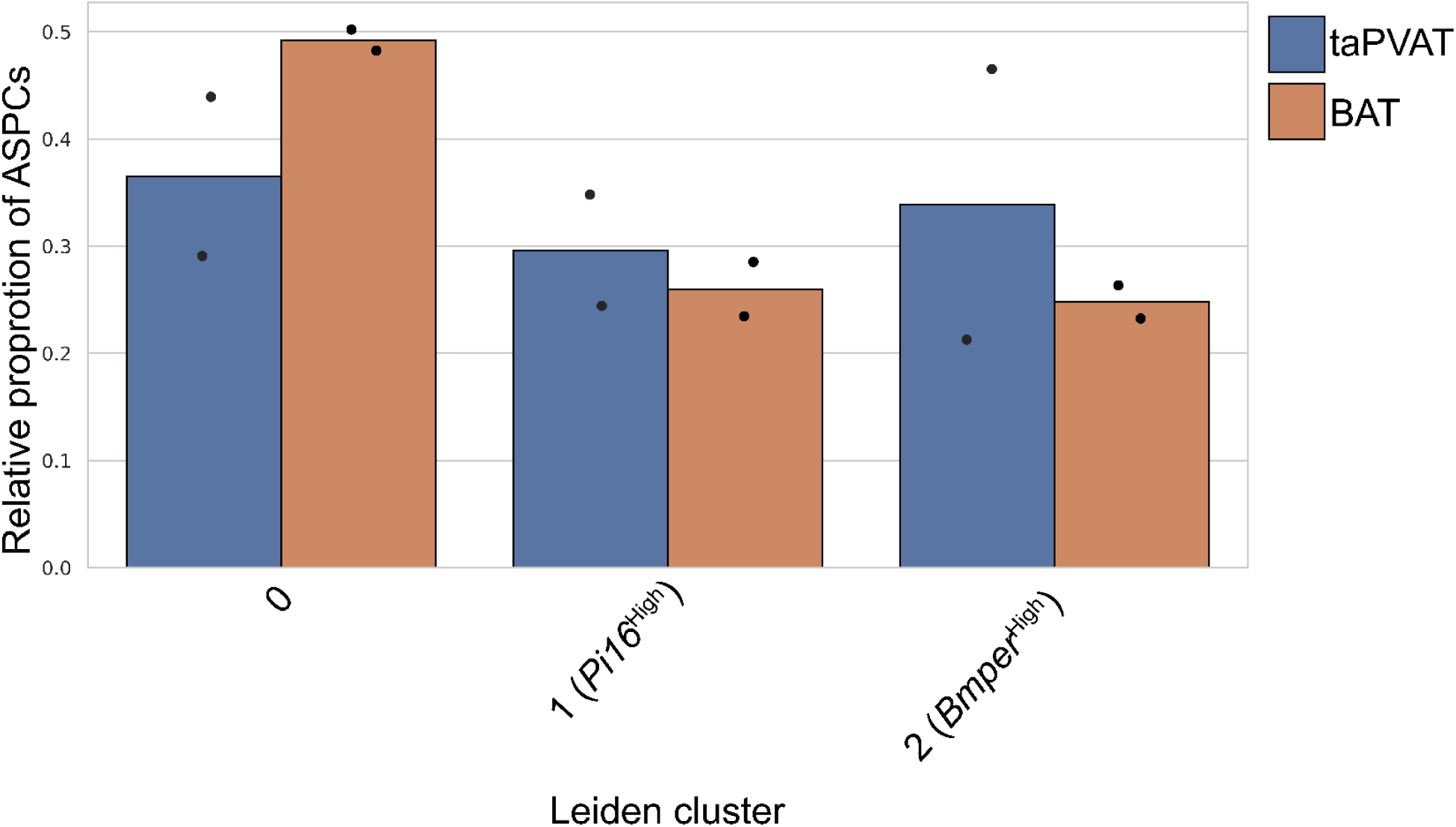
ASPC subpopulation proportions. The ASPC cluster was extracted from the full snRNAseq dataset, reintegrated, and reclustered to identify putative sub-populations. Three sub-populations were identified, each present in similar proportions. Bars represent the mean proportion for taPVAT and BAT and individual points show relative proportions in individual samples.

